# Taming the chaos gently: a Predictive Alignment learning rule in recurrent neural networks

**DOI:** 10.1101/2024.07.14.603423

**Authors:** Toshitake Asabuki, Claudia Clopath

## Abstract

Recurrent neural circuits often face inherent complexities in learning and generating their desired outputs, especially when they initially exhibit chaotic spontaneous activity. While the celebrated FORCE learning rule can train chaotic recurrent networks to produce coherent patterns by suppressing chaos, it requires non-local plasticity rules and extremely quick plasticity, raising the question of how synapses adapt on local, biologically plausible timescales to handle potential chaotic dynamics. We propose a novel framework called “Predictive Alignment”, which tames the chaotic recurrent dynamics to generate a variety of patterned activities via a biologically plausible plasticity rule. Unlike most recurrent learning rules, predictive alignment does not aim to directly minimize output error to train recurrent connections, but rather it tries to efficiently suppress chaos by aligning recurrent prediction with chaotic activity. We show that the proposed learning rule can perform supervised learning of multiple target signals, including complex low-dimensional attractors, delay matching tasks that require short-term temporal memory, and finally even dynamic movie clips with high-dimensional pixels. Our findings shed light on how predictions in recurrent circuits can support learning.

## Introduction

Humans and animals exhibit remarkable capabilities in learning and generating complex behaviors essential for a wide range of tasks. Indeed, the brain can accurately acquire and recall complex sequences, from the order of words in a sentence to highly skilled motor behaviors. The brain’s capacity to flexibly process sequential data and generate complex outputs is underpinned by the continuous, coordinated activity of neural networks, which integrate temporal and spatial information to precisely control its outputs.

Recurrent Neural Networks (RNNs) are powerful computational models that can capture and process sequential information while exhibiting complex dynamics (Rumelhart et al., 1986; Williams and Zipser, 1989; Pearlmutter, 1989; Atiya and Parlos, 2000). Cortical circuits often exhibit chaotic spontaneous activity (van Vreeswijk and Sompolinsky, 1996; Amit and Brunel, 1997; Brunel, 2000; Toyoizumi and Abbott, 2011; Rajan et al., 2016). While this chaotic behavior provides a variety of basis functions for network dynamics, the question arises as to how it can be transformed into desired patterned dynamics. The FORCE learning, which originated from reservoir computation (Lukoševičius et al., 2012; Lukoševičius and Jaeger, 2009; Maass et al., 2002; Jaeger, 2001; Ozturk and Principe, 2005; Maass et al., 2004; Wojcik and Kaminski, 2004), has been widely used as an algorithm for learning networks that exhibit chaotic behavior (Sussillo and Abbott, 2009). However, this approach relies on non-local weight updates with extremely fast synaptic changes, which seems to lack biological plausibility. How biologically plausible learning algorithms for recurrent neural networks can effectively leverage their rich dynamical properties to efficiently learn and recall complex sequential information remains an elusive challenge (Gilra and Gerstner, 2017; Murray, 2019; Bellec et al., 2020), especially when the networks show chaotic dynamics prior to learning.

In this paper, we introduce “Predictive Alignment”, an alternative learning framework designed to train recurrent neural networks over a variety of complex tasks while overcoming the limitations of existing methods. Our proposed learning rule modifies plastic recurrent connections to predict output feedback signals, while aligning these predictive dynamics with existing chaotic spontaneous dynamics (arising from fixed recurrent connections), which in turn suppress the chaos efficiently and improving network performance. The key innovation of predictive alignment lies in its ability to perform online and local supervised learning through prediction, enabling the network to learn multiple target signals efficiently and robustly. We demonstrate that Predictive Alignment can successfully train networks to generate diverse complex target signals with nonlinear dynamics, such as the chaotic Lorenz attractor, delay-matching tasks that require short term memory of temporal information, and high-dimensional spatiotemporal patterns in a movie clip. The proposed learning rule not only sheds light on how predictions can guide learning in circuits, but also provides a biologically plausible solution for training powerful recurrent networks in various applications.

## Results

### Recurrent neural network

We considered a rate-based recurrent network (Figure 1) with the following dynamics:

**Figure 1.**
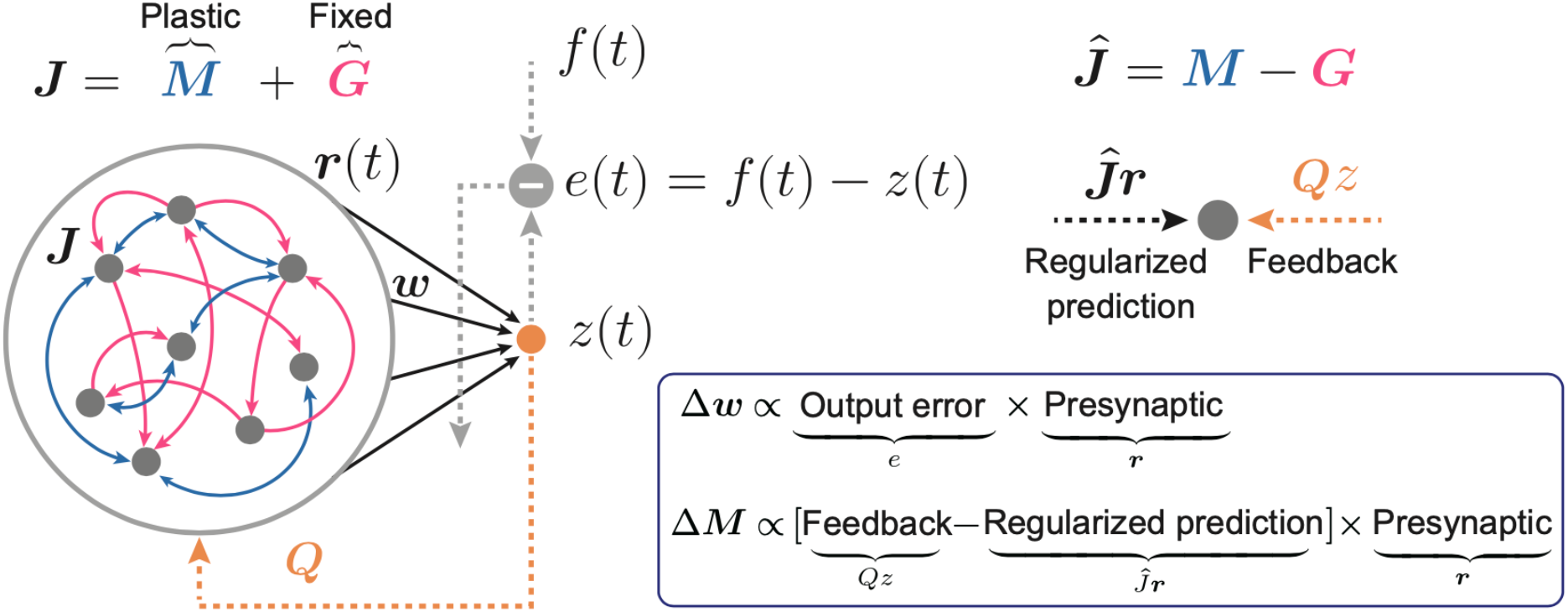
Recurrent neural network and predictive alignment rule. The network consists of recurrent layer and a readout unit. Multiple readouts will be considered later, yet only a single readout is illustrated in this example. Readout weight vector ***w*** is trained to minimize the error between a target and the readout activity (left figure, gray dashed arrows). Recurrent connection ***J*** is a summation of plastic yet initially weak connections ***M***, and strong and fixed connectivity ***G***. The plastic recurrent connections ***M*** is trained to minimize the error between feedback from output through random weights **Q** and a recurrent prediction ***Mr***, while aligning predictive recurrent dynamics to the chaotic dynamics.

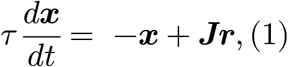

where ***x*** is a vector of membrane potentials of network neurons and *τ* is a time constant. The variable ***r*** is a firing rate vector, defined as ***r*** = tanh(***x***). The matrix ***J*** is a recurrent connectivity and is assumed to be a summation of two types of matrixes, ***M*** and ***G***:

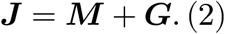

Although both types of connections are assumed to be initially generated with Gaussian distributions, they have some division of labor. First, we assumed that M is a plastic but initially weak connection, while G is a strong and fixed connection of which the large elements lead to chaotic network activity prior to learning (Sompolinsky et al., 1988; Sussillo and Abbott, 2009) (Methods). The recurrent plasticity rule we propose in this paper is applied to M, while G is always static. Second, these connections are assumed to have different sparseness: M has full connections while G has a sparse connection (Methods). After learning, M will suppress the chaotic spontaneous activity.

The output of network, or a “readout” is assumed to be a weighted linear summation of network activities:

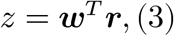

where ***w*** is a readout weight vector and *T* is a transpose. Multiple readouts can be defined, each with its own set of weight vectors.

### The Predictive Alignment learning rule

Similar to standard gradient descent rules, readout weights were trained to minimize the error between the target signals *f* and the model’s output *z* :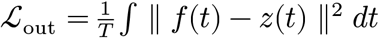, where *T* is a duration of learning phase. As we are considering online learning rule, at each time step, the resulting weight update rule can be written as:

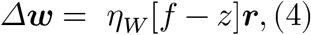

where *η*_*W*_ is a learning rate. For simplicity, the time dependence notation has been removed. The above rule for the readout weights is a standard least-meansquare (LMS) or also called the Delta-rule, hence not novel.

Predictive Alignment is a new learning rule for the recurrent weights. Unlike most learning rules for recurrent networks, which minimize the same cost function for the outputs (i.e., *ℒ*_out_ defined above), the aim of Predictive Alignment is to predict a feedback signal from the readout unit with the recurrent dynamics, while aligning predictive recurrent dynamics to the chaotic dynamics. More precisely, the plastic recurrent connectivity *M* is asked to minimize the cost function shown below:

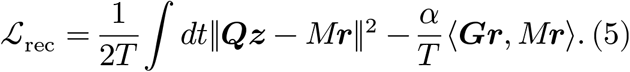

Here, the first term of the cost function is a deviation between the recurrent dynamics *M****r*** and the feedback signal ***Qz***, where ***Q*** is a static feedback connection. It should be noted that feedback from readout appears only in the learning rule, hence does not affect network dynamics directly. Minimizing this first term in the cost function requires the recurrent dynamics to predict the feedback signal. The second term is a regularization term, which plays a crucial role to suppress the chaos efficiently by aligning the predictive recurrent dynamics *M****r*** to the chaotic dynamics ***Gr***. We will show the effect of regularization in detail later. Unless other specified, we will assume *α* = 1 bthroughout the paper. The resulting online learning rule which minimizes the cost function *ℒ*_rec_ can then be written as:

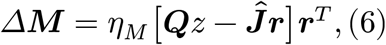

where we defined a regularized recurrent prediction as ***Ĵ***_*r*_ with ***Ĵ*** = ***M*** − *α****G*** (Methods).

In summary, we have proposed a set of plasticity rules for training chaotic recurrent neural networks. The readout weights are simply modified to minimize the error between the actual output and the target signal, while the recurrent connections are modified to minimize the feedback prediction error while aligning the predictive recurrent and chaotic dynamics – hence the name Predictive Alignment.

### Toy examples of learning with the Predictive Alignment rule

We first show some simple toy examples to demonstrate that Predictive Alignment can learn various forms of desired target signals. For the sake of simplicity, we assume periodic target signals in the toy examples, but we will consider non-periodic targets later. During the early phase of learning (Fig. 2A, 0-0.6 s), the readout activity (Fig. 2A, red trace) showed a mismatch from the target signal (Fig. 2A, green trace), indicating that the readout activity did not follow the target signal at this early stage. This contrasts with the behavior observed in the FORCE learning paradigm, which consistently clamps the output to the target through recursive least squares (RLS). As learning progressed, the readout error decreased monotonically, and the readout activity became similar to the target function (Fig. 2C). Once sufficient learning had occurred, the readout activity showed dynamics that were well matched to the target signal, even when plasticity was turned off (Figs. 2A, right side of vertical dotted line). Furthermore, while the recurrent neurons initially showed chaotic behavior, the activities became coherent and structured after sufficient learning (Figure 2B).

**Figure 2.**
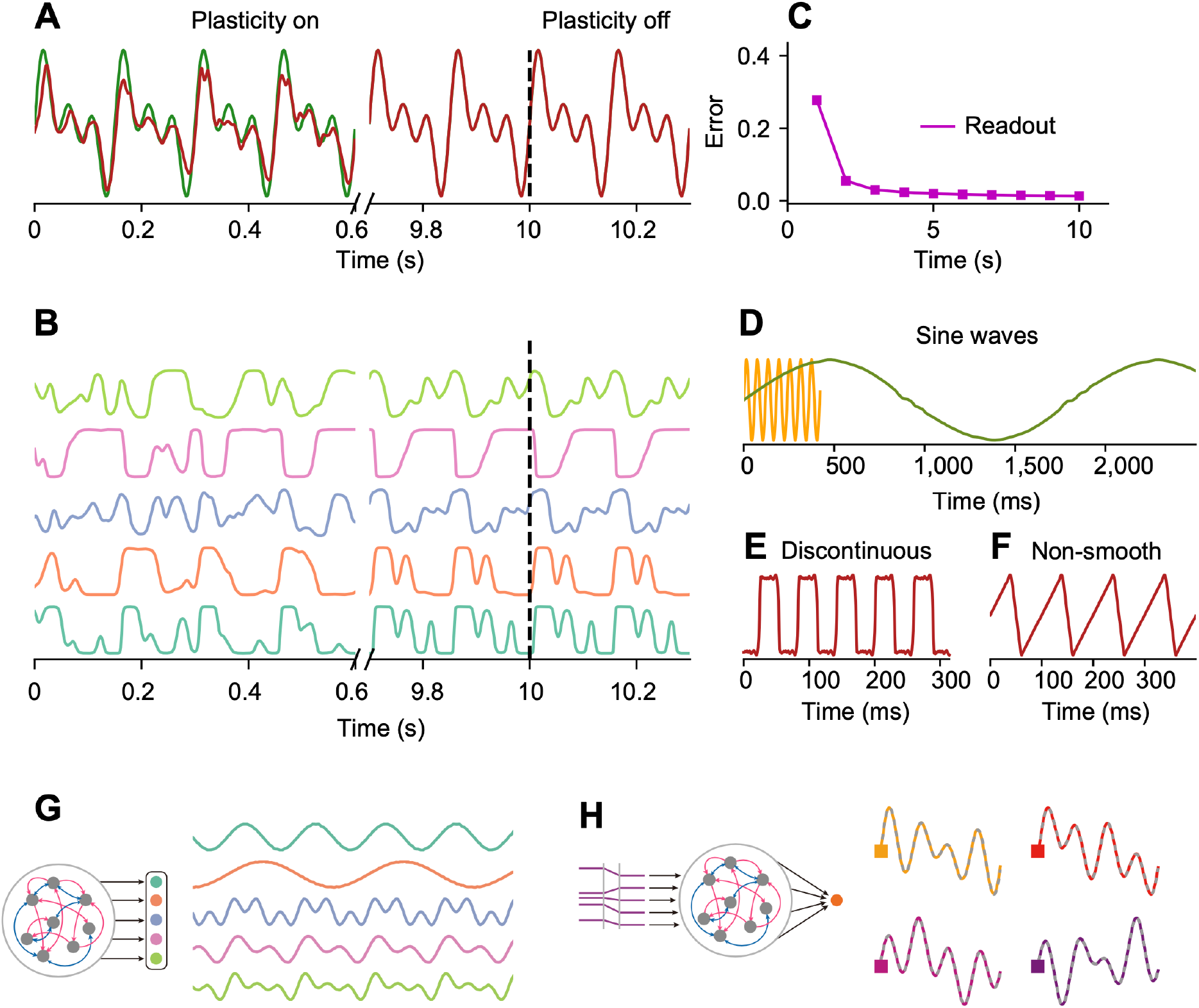
Examples of Predictive Alignment. (A) During the early phase of learning, the model output (red) deviated from the desired target (green), while it matched the output after sufficient training even after the plasticity was turned off. (B) The network activities initially showed chaotic activities and then was transformed to coherent activities after training. (C) Root mean squared error between the target and the output during training. (D-F) Examples of output activities trained to generate various simple target patterns. (D) The network learned sine waves *y* = 1.5sin (2*πt*/*T*)with different frequencies (*T* = 6*τ*; orange and *T* = 200*τ*; green). (E) The model learned discontinuous step-like target. (F) The model learned non-smooth sawtooth pattern. (G) A network with five readout units was considered (left). Such network could generate five distinct target patterns simultaneously (right). (H) A network receiving external control inputs (left) generates corresponding output patterns (right; colored lines). The corresponding target signals are shown as gray dashed lines superimposed on the targets.

Next, we examined how learning in the network changes the structure of the eigenvalue spectrum of the recurrent weight matrix. Before learning, the eigenvalues were uniformly distributed in a circle in the complex plane because the initial connectivity was a random matrix (Supplementary Fig. 1, left). After learning, while the connections had most of their eigenvalues within a circle in the complex plane, some pairs of leading eigenvalues with large real parts appeared (Supplementary Fig. 1, right). Such outliers in the spectrum generated by low-dimensional perturbations change the dynamics (Tao, 2013; Mastrogiuseppe and Ostojic, 2018).

We further show that the Predictive Alignment can learn more complex and diverse target patterns. The network can learn sinusoids with small (Fig. 2D, orange) or much larger (Fig. 2D, green) periods, as well as discontinuous (Figure 2E) and even non-smooth (Figure 2F) target signals.

In the above examples, we have demonstrated the performance of the network using simple examples. In particular, the network architecture considered above had only a single readout unit and learns only a single target signal. To see whether the Predictive Alignment still works for multiple targets, we first extended the learning paradigm beyond a single readout unit and make it face complex scenarios involving multiple readout units and corresponding target signals. Here, each recurrent neuron’s plasticity was modulated by a weighted combination of all output signals (Methods). In our simulation, we considered five periodic target signals, each was presented to one of readout units as a corresponding target signal. We found that the trained model exhibits five output patterns, distinctly aligned with the specified targets. This result shows the network’s ability to learn and generate diverse output patterns, without requiring that feedback signals are constrained to subpopulation-specific interactions; in our simulation, they indeed were all-to-all interactions across the entire recurrent population.

Having demonstrated that a network could learn multiple target signals, we wondered whether the network could perform input-output transformation tasks. To test this, we introduced four static control input patterns to the network neurons (Fig. 2H). Each desired output function was paired with a corresponding input pattern. The inputs were randomly assigned constant values, without temporal information. They acted only as switches to generate specific output functions. We found that such a network with a single readout unit could learn to generate different outputs depending on the given control input patterns (Fig. 2H).

Next, to confirm the robustness of the model, we fed the network with different strengths of noise while training the patterned target signal. We confirmed that while the model’s output error increased as the noise intensity was increased, the degree of increase was sufficiently small compared to FORCE (Supplementary Fig. 2).

In summary, we have tested our Predictive Alignment learning rule over various forms of target signals and multiple readouts. Further, the model can transform static input signals to the transient output patterns by modifying the recurrent connectivity.

### Recurrent prediction aligns to the chaotic dynamics

To understand the mechanism underlying the Predictive Alignment, we next analyzed the effect of recurrent plasticity in the network undergoing learning of a simple periodic signal (as shown in Figure 2A). Learning of the recurrent connections minimizes of error between the regularized recurrent dynamics and the feedback signal (Equation 6). The error therefore dropped monotonically and a plastic recurrent dynamics (***Ĵr***) gradually matched to the feedback signal (***Q****z*) (Fig. 3A, B). We would emphasize that the term ***Ĵr*** is the regularized recurrent prediction (see Eq. 6). Recall that minimization of our cost function (Eq. 5) requires alignment of the predictive recurrent dynamics ***Mr*** to the chaotic dynamics ***Gr***. In the following, we will call such an alignment of two types of dynamics “recurrent alignment”.

**Figure 3.**
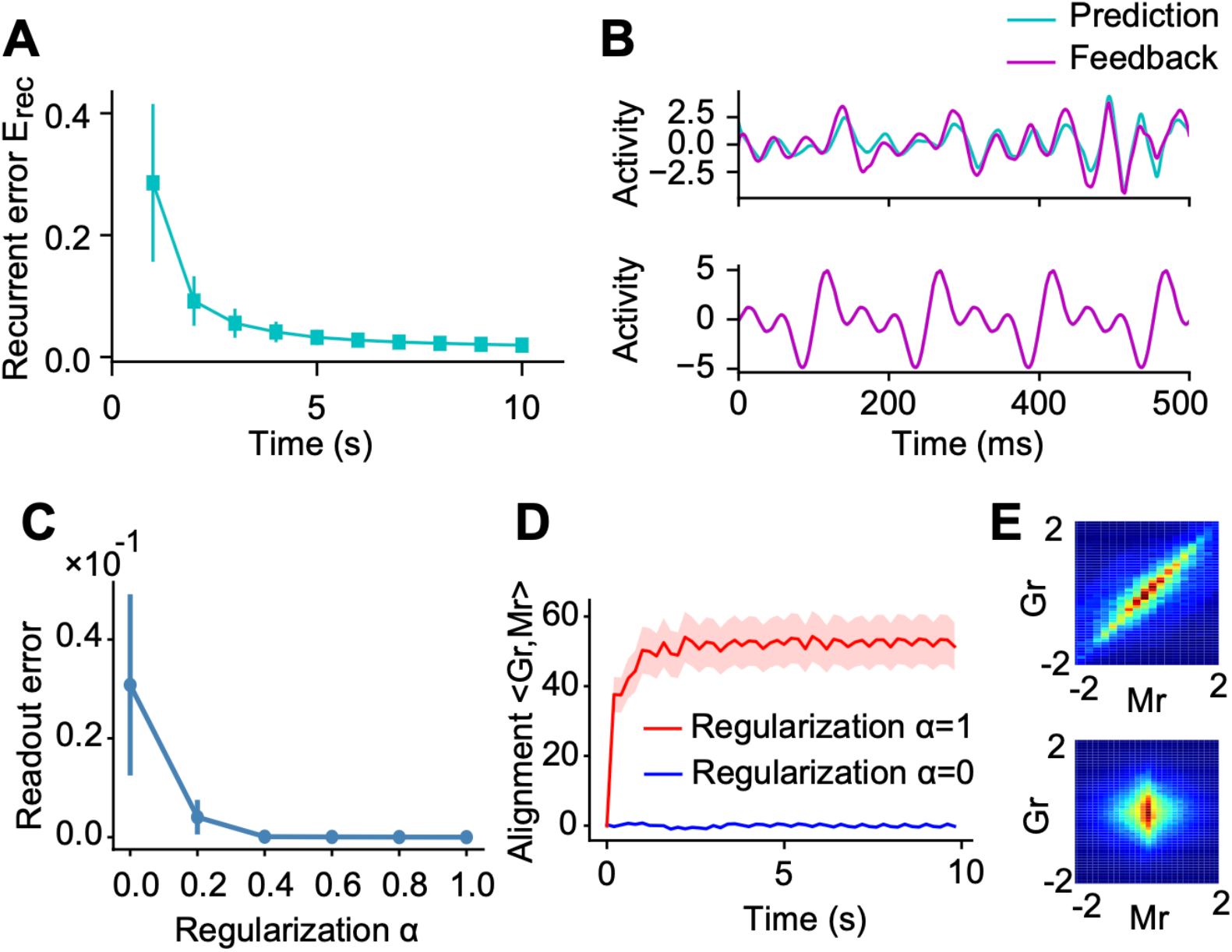
Network mechanism of predictive alignment. (A) The recurrent error 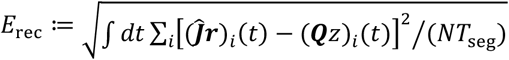 between the regularized recurrent prediction ***Ĵr*** and the feedback signal ***Q****z* during training. Here, *N* is the number of neurons in the network and *T*_seg_ denotes the duration of each time segment, with the entire time period equally divided into 10 segments. The time integration was performed over the duration of each segment. (B) Example dynamics of the regularized recurrent prediction ***Ĵr*** and the feedback signal ***Q****z* during early (top) and late (bottom) phases of learning are shown. (C) Root mean squared errors between the target and the trained output over different values of the regularization parameter are shown. (D) The dynamics of the correlation between the plastic recurrent *M****r*** and the chaotic ***Gr*** dynamics are shown for with (red) and without (blue) regularization. (E) Joint distributions of plastic recurrent *M****r*** and chaotic ***Gr*** dynamics are shown with (top) and without (bottom) regularization. In A and C, error bars stand for s.d.s over 20 independent simulations. In D, shaded areas represent s.d.s over 10 independent simulations.

We next demonstrate the role of recurrent alignment in our plasticity rule. To this end, we first calculated the output error over various degree of regularization parameter *α*. We found that increasing the value of α resulted in a reduction in the model output error relative to the target function (Fig. 3C). Interestingly, the variance of the error over multiple simulations also decreased with higher values of α. These results suggest that larger α increases the accuracy and stability of learning. To further understand the mechanism underlying these results, we first calculated the correlation term in the cost function (i.e., ⟨***Gr***, *M****r***⟩) over the entire learning period in two cases. For the first case, we trained the network with *α* = 1, indicating that the learning rule reduces the error under a trade-off with the regularization term (aligned case). In contrast, in the second case, the recurrent connections were trained with *α* = 0, hence such trade-off was not considered (control case). As expected, the value of correlation grew in the aligned case, while such behavior was not observed in the control case (Fig. 3D, E).

Altogether, these results indicate that the Predictive Alignment suppresses the chaotic activity more efficiently by aligning the recurrent prediction to the chaotic dynamics, allowing for robust computation.

### Learning performance around the edge of chaos

While we have introduced a static strong recurrent connectivity G, the role of spontaneous chaotic activity generated by such strong connectivity remains to be explored. Following the standard rate-based recurrent network, the strength of static connectivity within the network was scaled by a factor g (Sompolinsky et al., 1988). Recurrent networks with g < 1 produce decaying activity in the spontaneous activity regime, while g > 1 leads to irregular chaotic spontaneous activity (Sompolinsky et al., 1988). In FORCE learning, initial networks with g just above such a critical point, the so-called “edge of chaos”, show the best performance for learning.

We wondered to what extent the performance of our model depends on the strength of the scaling factor g. To test this, we first measured the average root mean square (RMS) error after training, across different networks with different values of g. Similar to FORCE, the error between the target function and the output of the network shows best performance when the scaling factor is just above 1.0 (i.e., the edge of chaos) (Supplementary Fig. 3A). We found that the RMS error between the recurrent prediction and the feedback also showed a minimum value when the scaling factor g was around the edge of chaos (Supplementary Fig. 3B). We then measured the strength of the output weight vector and the recurrent connection matrix. Large values of the weights can lead to training instability and also make the network more sensitive to noise, indicating that the network is losing robustness. We found that both weights had a minimum when the network is on the edge of chaos initially (Supplementary Fig. 3C, D).

Taken together, these results suggest that Predictive Alignment produces accurate and robust output when the initial network state is on the edge of chaos. This chaos is then tamed slowly by the recurrent alignment (see Fig. 3).

### Learning low-dimensional chaotic attractor: the Lorenz attractor

The tasks we have shown so far are relatively simple, with periodic targets only. This raises the question of whether our models can learn more complex dynamics such as the three-dimensional Lorenz attractor (shown in Figure 4G) as a target trajectory. Unlike the simple periodic signals studied in the previous scenarios, the Lorenz attractor exhibits non-periodic and complex behavior.

**Figure 4.**
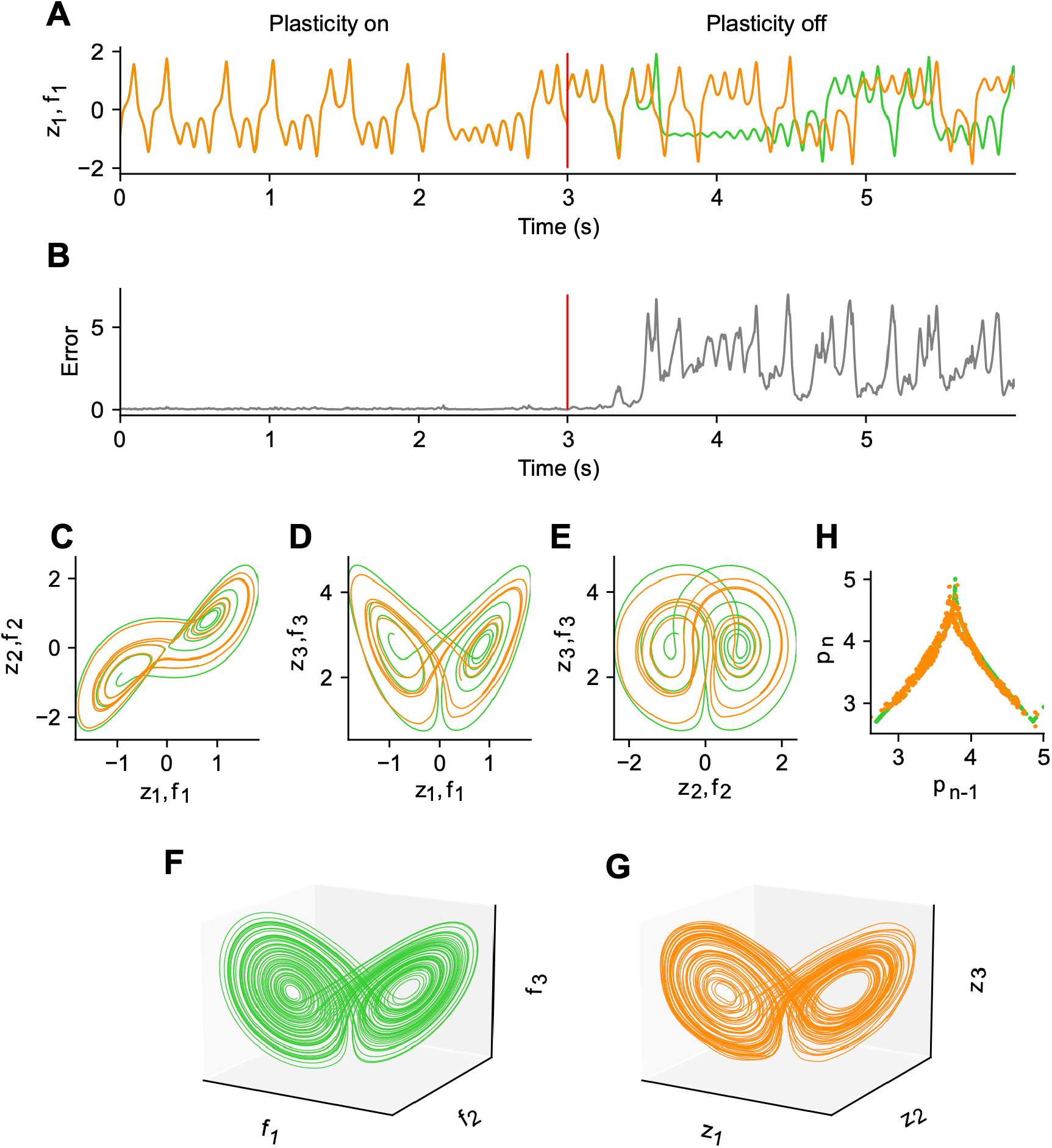
Learning chaotic dynamics: the Lorenz attractor. (A) First component of target Lorenz attractor signal (green) and the output activity of the network (orange). The vertical red line represents the time when plasticity was turned off from on after learning. The output of the network after training showed almost the same dynamics as the target signal when the plasticity was on. Even after the plasticity was turned off, the output initially showed the same dynamics as the target, but after a while it showed a pattern different from the target but similar to Lorenz dynamics. (B) Error between the two signals shown in A is shown. (C-E) Trajectories projected on two-dimensional state spaces are shown. (F) Target Lorenz attractor in three-dimensional state space is shown. (G) Same as in F, but for the outputs. (H) Tent map representation plotting the current local maximum of the third component against the previous local maximum for the target signal (green) and network output (orange) is shown.

We considered a network with three readout units to learn the three-dimensional Lorenz attractor. Even though the desired targets are not periodic, the late phase of learning showed readout dynamics that closely matched the desired output (Fig. 4A, 0-3 s). We found that, when plasticity was off, the output predicted the subsequent dynamics well, but then showed divergence from the targets because the model itself is a chaotic system. Interestingly, despite this deviation from the desired dynamics, the readout activity still showed complex oscillations similar to those of the target.

We then compared the trajectories of the dynamics of the trained network when plasticity was off and the target signals. Both the projected trajectories on the twodimensional planes and the full trajectories show striking similarities between the two, suggesting that the model learns the manifold representative of the true Lorenz attractor (Figs. 4C-G). Quantitative analysis of the tent map (Lorenz, 1963) of successive maxima relative to the previous maxima further indicates that the model has adequately learned the Lorenz attractor (Fig. 4H).

In summary, we have shown that the Predictive Alignment can learn and generate low-dimensional complex autonomous dynamics through recurrent plasticity.

### Learning generalized representations

The above results have demonstrated that the Predictive Alignment can learn and generate complex target signals by adapting both readout and recurrent connections through plasticity. This was achieved by precise adjustment of both recurrent and output weights. In a machine learning framework called Reservoir Computing (RC), it has been shown that recurrent networks can learn complex tasks without relying on the plasticity of recurrent connections (Lukoševičius et al., 2012; Lukoševičius and Jaeger, 2009; Maass et al., 2002; Jaeger, 2001; Ozturk and Principe, 2005; Maass et al., 2004; Wojcik and Kaminski, 2004). This is achieved by generating rich basis function in the network dynamics such that the readout unit can decode the arbitrary output signals. Inspired by such a rich generalization ability in the recurrent networks, we then explored the model’s potential to generalize pretrained simpler output patterns to more complex targets without recurrent plasticity.

We considered a structured task with a network consisting of two different groups of readout units to answer our above question. The first group underwent the Predictive Alignment learning to generate a set of simple multi-frequency sinusoidal signals (Supplementary Fig. 4A; multiple sinusoids). During this phase, both recurrent and readout weights were trained on the first group of readout units, while the weights on the second group of readouts remained static. Once the first group was successfully trained to generate sinusoidal targets, we introduced a second phase into the learning process, in which recurrent plasticity was terminated while the readout weights of the second group were started to train to generate novel target signals (Supplementary Fig. 4B). We basically used our pretrained network with the Predictive Alignment as a reservoir computing network. Note that, in principle, any periodic function can be approximated by a summation of a sufficient number of sinusoidal components. We found that the second group successfully learned and generated novel signals, even though the recurrent connections were not trained to optimize for such targets. In contrast, the network without training in the first phase failed to learn the target signals in the second phase, as the network did not learn the sinusoids (Supplementary Figs. 4C, D).

In summary, we have shown that the network can generalize from learning simple multi-frequency oscillations to learn and generate novel signals without relying on the plasticity of recurrent connections.

### Learning delay-matching task

Up to now, we have explored whether the network can learn autonomous dynamics or input-output transformations. We then wondered whether the proposed mechanism could learn more complex task, Measure-Wait-Go (MWG), in which the desired output was indicated by the time interval between two identical pulse inputs. In the MWG task, the network was required to measure that interval, keep it in memory during a delay period of random duration, and then reproduce it after the set signal (Jazayeri and Shadlen, 2010). This task is more difficult than the production of a periodic output due to the requirement for the RNN to learn to store the information about the interpulse delay, and then produce responses at different times depending on the value of delay.

To mimic such an MWG in our simulation, we considered two input neurons sending pulses to the network. In each trial, one of four delay values T_delay_ was sampled, and the two input neurons generated pulses with an interpulse delay of T_delay_ (Fig. 5A). The network projected to a single readout unit and the synapses were modified such that the output should be a pulse delayed by T_delay_ relative to the second input pulse. After learning, network generated the output with the desired time delay over all four samples (Fig. 5B, colored squares). More interestingly, the network interpolated well to input intervals in between those used for training, while failed to extrapolation (Fig. 5B, gray curve; Fig. 5C). These results suggest that the network did not simply learn individual output mappings, but instead learned an underlying manifold structure encoding the range of possible delays.

**Figure 5.**
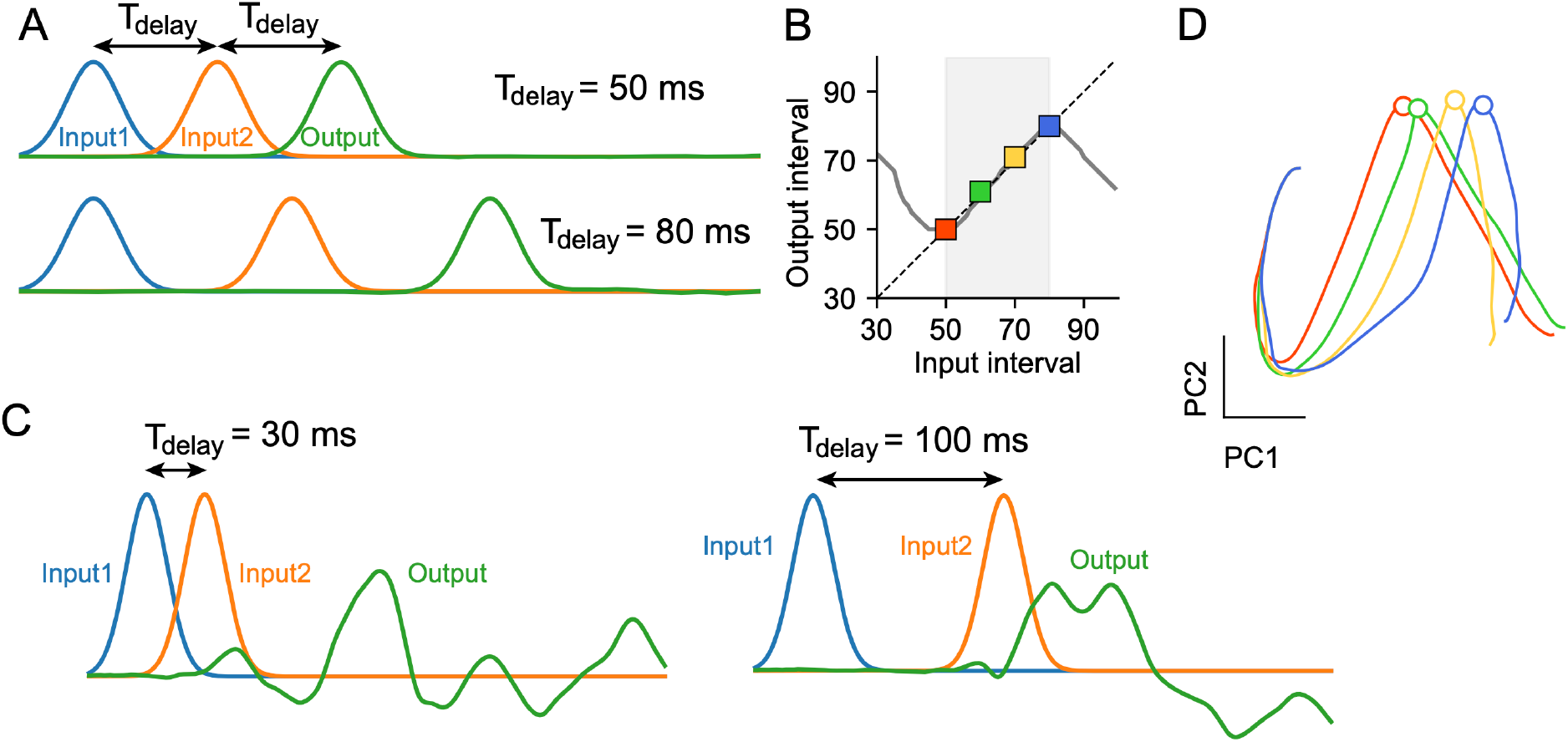
Learning delay-matching task. (A) In each trial, two input neurons (blue and orange) send pulses to the network with a random delay T_delay_ between pulses. The target network output (green) should be a pulse delayed by T_delay_ relative to the second input pulse. (B) Networks were trained on a set of samples (colored squares) and tested for generalization to novel inputs, including interpolation within the training range (shaded region) as well as extrapolation beyond the training range. (C) The network after training failed to extrapolate beyond the training range. (D) Principal component analysis (PCA) of the trained network revealed that as the delay T_delay_ increased, the network states corresponding to the output peak shifted linearly along a manifold in state space.

We then wondered whether the low dimensional network dynamics already shows the structured representations for the temporal delay information in the task. To test this prediction, we performed the principal component analysis (PCA) on the trained network and visualized the low dimensional representation in the network. Projected dynamics on the two-dimensional PC axis showed linear shifts of trajectories along the linear manifold with increasing delay, revealing that the temporal structure of the task is embedded in the lowdimensional recurrent dynamics (Fig. 5D). Taken together, these results demonstrate the network’s ability to generalize by acquiring the underlying manifold structure inherent to the task.

### Learning and replay of high-dimensional movie data

Finally, we demonstrate that the model can learn natural high-dimensional signals. Here, we train recurrent neural to learn and replay high-dimensional video patterns in pixel space. The original video sequence consisted of 100 frames. However, to enable learning of finer temporal dynamics, this sequence was temporally upsampled by interpolation to a higher resolution of 1,000 frames for use during training. Each individual video frame had a spatial dimension of 80×92 pixels, with three color channels (RGB) per pixel, resulting in a total of 22,080 units representing the pixel values (Fig. 6A). In contrast to the extremely highdimensional target signals, the recurrent neural network model itself consisted of only 800 neuron units. During training, synapses were modified to minimize the error between 22,080 readout activities and the movie data.

**Figure 6.**
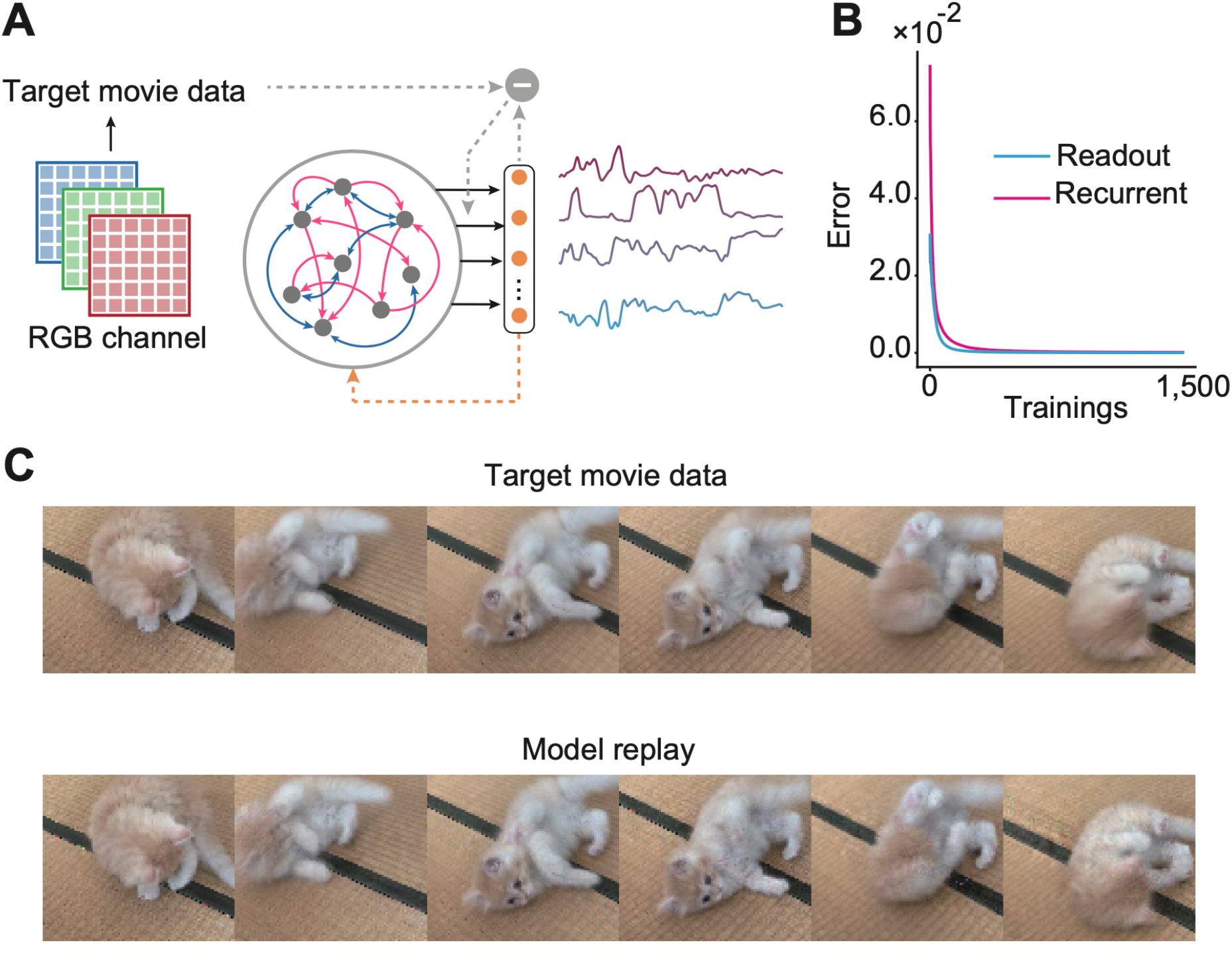
Predictive Alignment for movie data storage and replay. (A) Schematic of the network architecture for learning video sequences, with 22,080 pixel target signals and 800 neuron units in the recurrent network. (B) Learning curves showing the monotonic decrease in readout weight error (cyan) and recurrent weight change (magenta) over 1,500 epochs of training on the upsampled 1,000-frame video sequence. (C) Samples of the original video frames (top) compared to the output frames generated by the network (bottom) after training. Full movie data is available in Supplementary Movie 1.

Despite the large dimensionality mismatch between the target signals and the recurrent layer, the readout and recurrent weight training errors decreased monotonically and converged to near zero values after sufficient training iterations, suggesting the network dynamics were able to be trained successfully (Fig. 6B). Indeed, readout activities driven by the autonomous network dynamics after learning showed that the network dynamics accurately encoded and replayed the full training video patterns (Fig. 6C; Supplementary Movie 1). These results indicate that the proposed Predictive Alignment is not restricted to few target signals but can be applicable to high dimensional signals.

## Discussion

Many cortical and subcortical circuits exhibit complex, dynamic patterns of activity that underlie crucial cognitive functions such as working memory (Lundqvist et al. 2016), decision-making (Kepecs et al. 2008; Runyan et al. 2017), motor control (Churchland et al. 2012), and sensory processing (Stringer et al. 2019). In this paper, we have presented a novel learning framework which we call “Predictive Alignment” that enables recurrent neural networks to learn and generate a variety of patterned activities, even when the network initially exhibits chaotic spontaneous dynamics. The key insight of our approach is that instead of directly minimizing the output error to train recurrent connections, as in most existing recurrent learning rules (Robinson and Fallside, 1987; Werbos, 1988; Williams and Zipser, 1989; Hochreiter et al., 1997; Sussillo and Abbott, 2009; Sussillo and Abbott, 2012), we instead focus on predicting the recurrent activity itself and aligning this prediction with the chaotic dynamics of the network. This allows us to efficiently suppress the chaotic spontaneous activity and shape the network dynamics to produce the desired patterns.

The Predictive Alignment learning rule we proposed has several advantages in terms of biological plausibility. First, the synaptic plasticity rules are local, depending only on the activity of the pre- and post-synaptic neurons and their predicted activities, without requiring non-local information such as the inverse of the correlation matrix of the network activities (Sussillo, D., and Abbott, 2009) or unfolding the dynamics through time (Robinson and Fallside, 1987; Werbos, 1988). Second, our framework does not rely on clamping the network’s outputs to target signals. FORCE learning and its variants (Sussillo and Abbott, 2009, 2012; DePasquale et al., 2016; Thalmeier et al., 2016; Nicola and Clopath, 2016) tame the chaos by clamping the output to be close to the target during learning. They use weight changes which are faster than the time scale of the dynamics, which seems to lack biological plausibility. Instead, our Predictive Alignment learns to generate the desired patterns by predicting the feedback signal with aligning the recurrent and the chaotic dynamics, which does not require such assumptions. Since our rule uses multiple localized activities within each neuron (e.g., *M****r*** and ***Gr***), we expect that multicompartmental recordings from individual neurons during a learning task would enable to test whether biological neural circuits use similar local predictive learning rules. The proposed rule predicts that the correlation between multiple dendritic synaptic currents and/or somatic membrane potentials should increase as learning progresses. Further experimental studies are needed to better understand how such rules might give rise to structured neural dynamics.

Recently, several local learning rules for recurrent neural networks have been proposed, such as FOLLOW (Gilra and Gerstner, 2017) and RFLO (Murray, 2019). While the FOLLOW rule relies on clamping outputs to target values similar to FORCE, it achieves this clamping via negative error feedback from an autoencoder, allowing slower weight changes that make the error signal locally available at post-synaptic neurons. In contrast, our model does not require clamping at all because it can learn to generate desired output patterns even when the initial recurrent dynamics differ from the target, as described above. Furthermore, because FOLLOW drives outputs based on input dynamics, it cannot perform the delayed adaptation tasks considered here (Murray, 2019). Another framework, RFLO, approximates backpropagating output errors through random feedback connections to train the recurrent connections (Murray, 2019). While both our Predictive Alignment and the RFLO rely on random feedback projection from the output, the two rules have significant difference: the RFLO feedback the output error signal, whereas our rule adapts recurrent predictions to the output signals directly without relying on output errors.

Another powerful learning framework that approximates Backpropagation Through Time (BPTT) (Robinson and Fallside, 1987; Werbos, 1988), which is the most celebrated training method for recurrent neural networks in machine learning, is the e-prop (Bellec et al., 2020). In the e-prop, the gradual changes in the hidden variables of neurons generate eligibility traces that extend over long periods of time which in turn are combined with subsequent instantaneous error signals. While the Predictive Alignment also combines the instantaneous error signal and the presynaptic activity, there are some differences between the two. First, although the e-prop assumes low-passed presynaptic activity as an eligibility trace, our rule considers instantaneous presynaptic activity. Second, our rule requires the instantaneous error between the output activity via the feedback signal ***Q****z* and the regularized recurrent prediction ***Ĵr***, whereas the e-prop utilizes the error between the desired target signal and the output activity. Future work should clarify whether the proposed rule can approximate BPTT in some way, as e-prop does.

How can the Predictive Alignment framework be realistically implemented in the brain? The motor cortex provides a prime example of a recurrent circuit capable of learning complex temporal patterns. Neurons in the motor cortex receive input about the planned action from the premotor cortex and information about the current state of the body from sensory input. They are thought to use this combined signal to compute the error necessary for learning and adjusting their neural activity patterns to generate the desired output trajectory (Wolpert and Ghahramani, 2000). In the Predictive Alignment framework, instead of transmitting an error signal derived from a target output, the feedback from the network’s own output could be transmitted to each neuron as a “teacher” signal. Previous experimental studies suggest that the apical dendritic compartment receives input from higher cortical layers, which in turn is attenuated before reaching the somatic compartment (Larkum et al., 2009). Our framework predicts that this attenuated apical input does not directly influence somatic firing rates, but instead acts as a teaching signal that incorporates predictive feedback, allowing individual neurons to modify their recurrent connections to better anticipate and generate the required motor outputs. Further experimental studies would be needed to understand the specific biological mechanisms that could organize such a predictive alignment process within recurrent cortical circuits.

We considered the Predictive Alignment in a recurrent network with balanced excitatory and inhibitory synaptic currents. The implementation we used in this paper does not satisfy Dale’s law (Strata and Harvery, 1999). A more biologically plausible model would require separate excitatory and inhibitory neuron populations (Ingrosso and Abbott, 2019). It would be an interesting question how maintaining an appropriate excitation/inhibition balance via inhibitory plasticity (Vogels et al., 2011; Zenke et al., 2015; Litwin-Kumar and Doiron, 2014; Agnes and Vogels, 2024) is critical for stable learning in such two-population network with chaotic spontaneous activity. Future extensions of our model should explore more biologically plausible architectures consists of spiking neurons as well (Abbott et al., 2016; Thalmeier et al., 2016; Boerlin et al., 2013; Schwemmer et al., 2015; Bourdoukan and Deneve., 2015; Eliasmith et al., 2012; Kim and Chow, 2018). Another interesting direction would be to explore how our predictive plasticity interacts with reward signals. Reward signals could potentially guide the shaping of top-down feedback signals (Tajima et al., 2017), with Predictive Alignment then fine-tuning the dynamics. Alternatively, predictive and reward-based plasticity could operate in parallel on distinct synaptic subpopulations. Combining these plasticity mechanisms could provide new insights how different forms of learning interact to give rise to robust and flexible neural computation in reinforcement learning.

In conclusion, the Predictive Alignment presents a novel and biologically plausible learning approach for training chaotic recurrent neural networks. This framework suggests that prediction within the local circuit can guide powerful, robust and generalizable learning.

## Methods

### The Predictive Alignment learning rule

Our network consists of *N* rate-based neurons, mutually connected with the recurrent connections. Here, we considered two types of recurrent connections: strong and fixed connectivity G and initially weak yet plastic connectivity M. The strength of each connection was generated by a Gaussian distribution, and unless other specified, with zero-mean and the standard deviation of 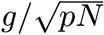, with (*p, g*) = (0.1, 1.2) for G and (*p, g*) = (1.0, 0.5) for M. Here, *g* is a gain of connection, leading chaotic spontaneous activity with *g* > 1, thus the fixed recurrent connectivity G generates initial chaotic spontaneous activity. The dynamics of membrane potential with the external input ***I*** was governed by the following equation:

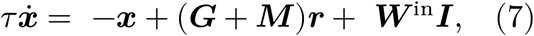

where ***x*** is a membrane potential and ***r*** is a firing rate of neuron, defined as:

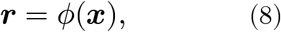

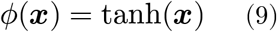

The matrix ***W***^in^ is a feedforward connectivity projecting from input layer, which is assumed to be fixed during the whole simulation. The parameter *τ* is a membrane time constant, which we set as *τ* = 10 (ms). The recurrent neurons project to readout neurons, of which the output value was defined as a linear summation of firing rates:

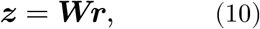

where ***W*** is a readout weight matrix.

The cost function for the plastic recurrent connectivity is defined over a period of learning phase *T* as follows:

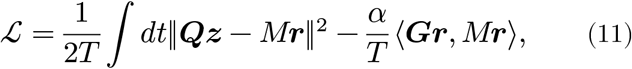

where ***Q*** ∈ ℝ^*N*×*K*^ is a random feedback matrix projecting from the readouts and ⟨***Gr***, *M****r***⟩ is a correlation defined as:

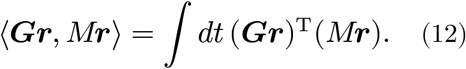

All elements of ***Q*** were chosen randomly and uniformly over the range 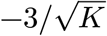 to 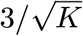, where *K* is a number of readouts. Taking the gradient of this cost function, we obtain an online learning rule as:

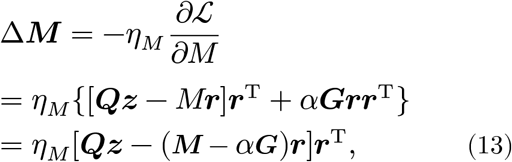

where T is a transpose. Readout weights were trained by simple least-meansquare (LMS):

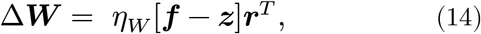

where ***f*** is a vector of teaching signal for the model’s outputs.

#### Learning Lorenz attractor

In Fig.4, we trained a network with three readout units with Lorenz attractor as the target signal following the dynamics below:

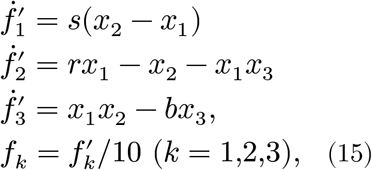

where (*s, r, b*) = (10, 28, 8/3) in our simulation. Learning was performed using a 15,000 second trajectory generated according to these dynamics as a target signal.

### Simulation details

All simulations were performed in customized Python3 code written by TA with numpy 1.17.3 and scipy 0.18. Differential equations were numerically integrated using a Euler method with integration time steps of 1 ms.

## Acknowledgments

The authors express their sincere thanks to Guillaume Bellec for his valuable comments on our manuscript. This work was supported by BBSRC BB/N013956/1, BB/N019008/1, Wellcome Trust 200790/Z/16/Z, Simons Foundation 564408 and EPSRC EP/R035806/1.

## Author Contributions

T.A. and C.C. conceived the study and wrote the paper.

T.A. performed the simulations and data analyses.

---

**Supplementary Figure 1.**
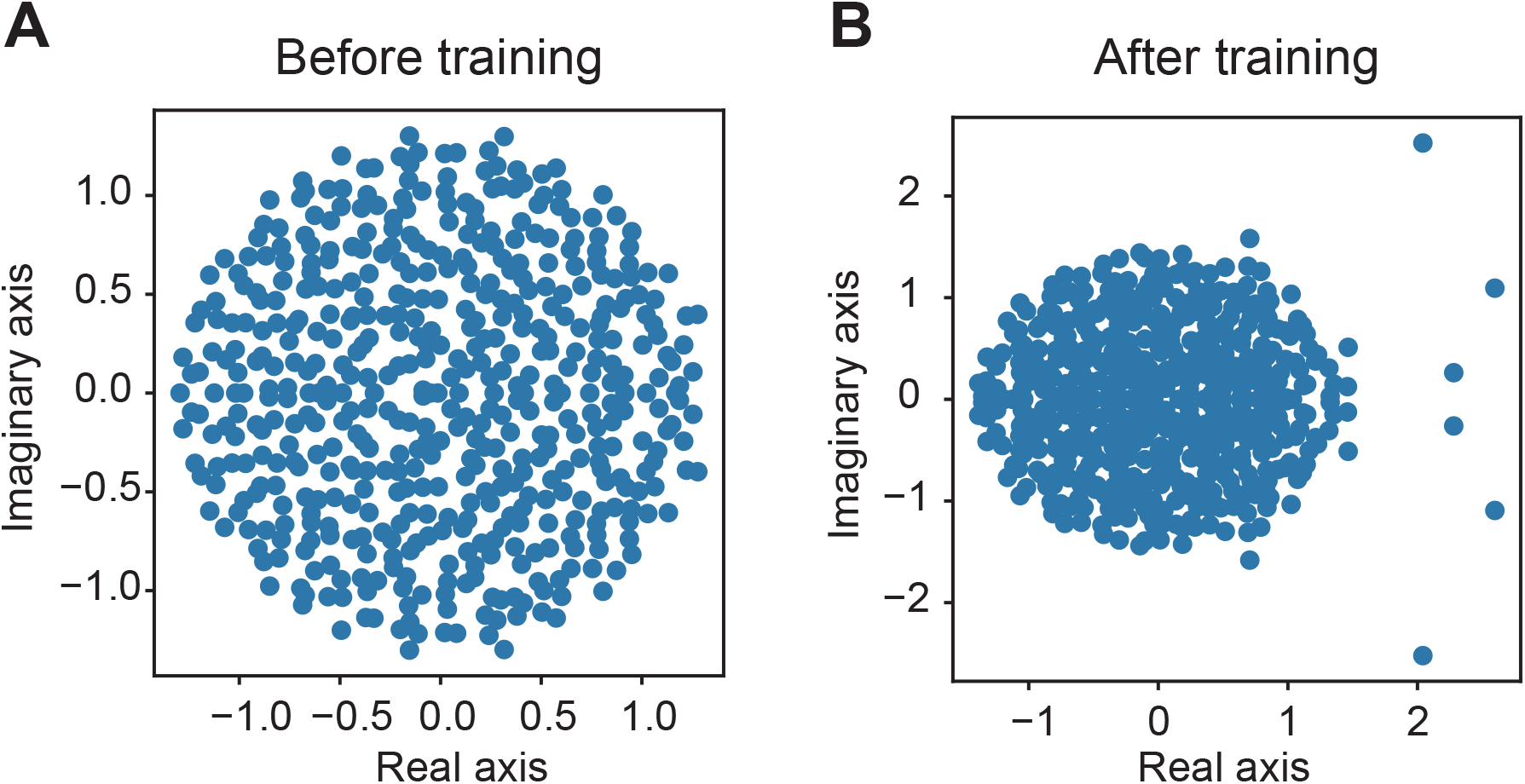
Eigenvalue spectrum of the recurrent connections. (A) The eigenvalues are uniformly distributed within a circle, since the initial recurrent connections were generated by Gaussian distribution. (B) The spectrum of weights after training shows that most eigenvalues still lie on a circle in the complex plane, while only a few eigenvalues lie outside the circle.

**Supplementary Figure 2.**
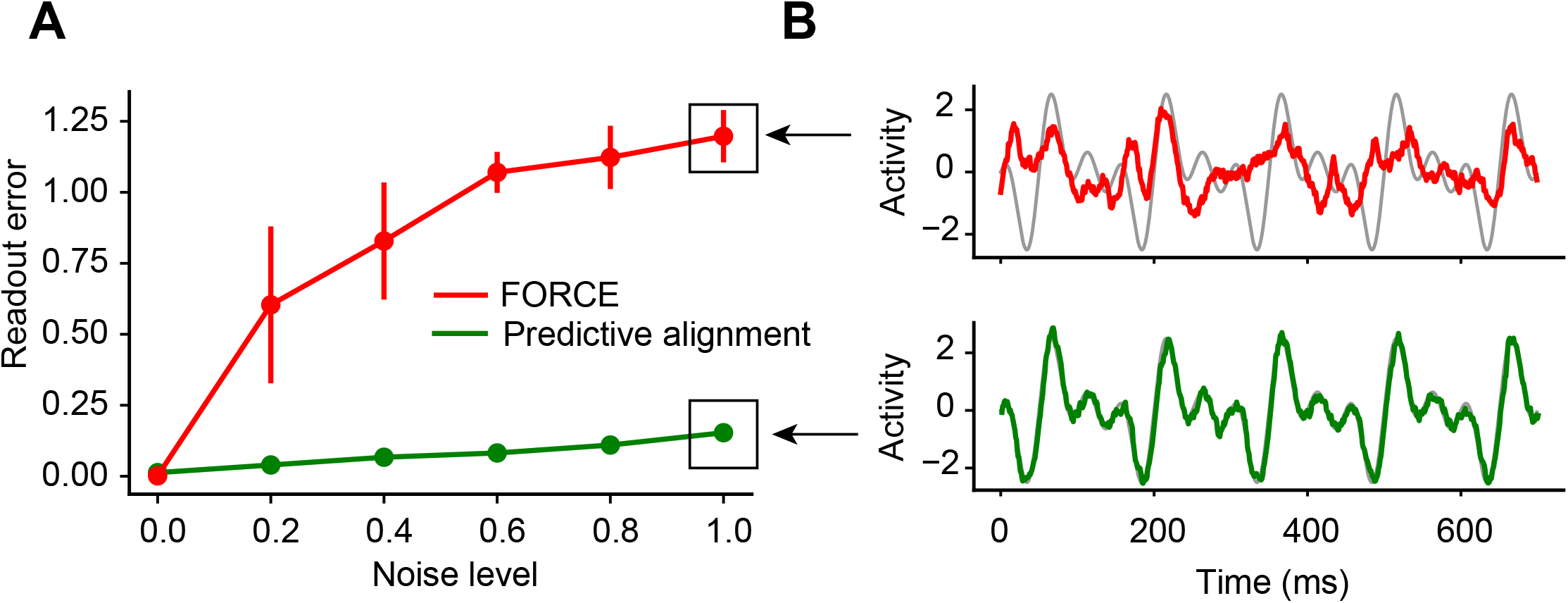
Robustness of predictive alignment against noise. (A) Networks with different strengths of noise were trained with the FORCE (red) and the predictive alignment (green) with the patterned target signal. Error bars stand for s.d.s over 20 independent simulations. (B) Example readout activities during the late phase of training are shown. Colors are the same as in A. The gray traces represent the target signal.

**Supplementary Figure 3.**
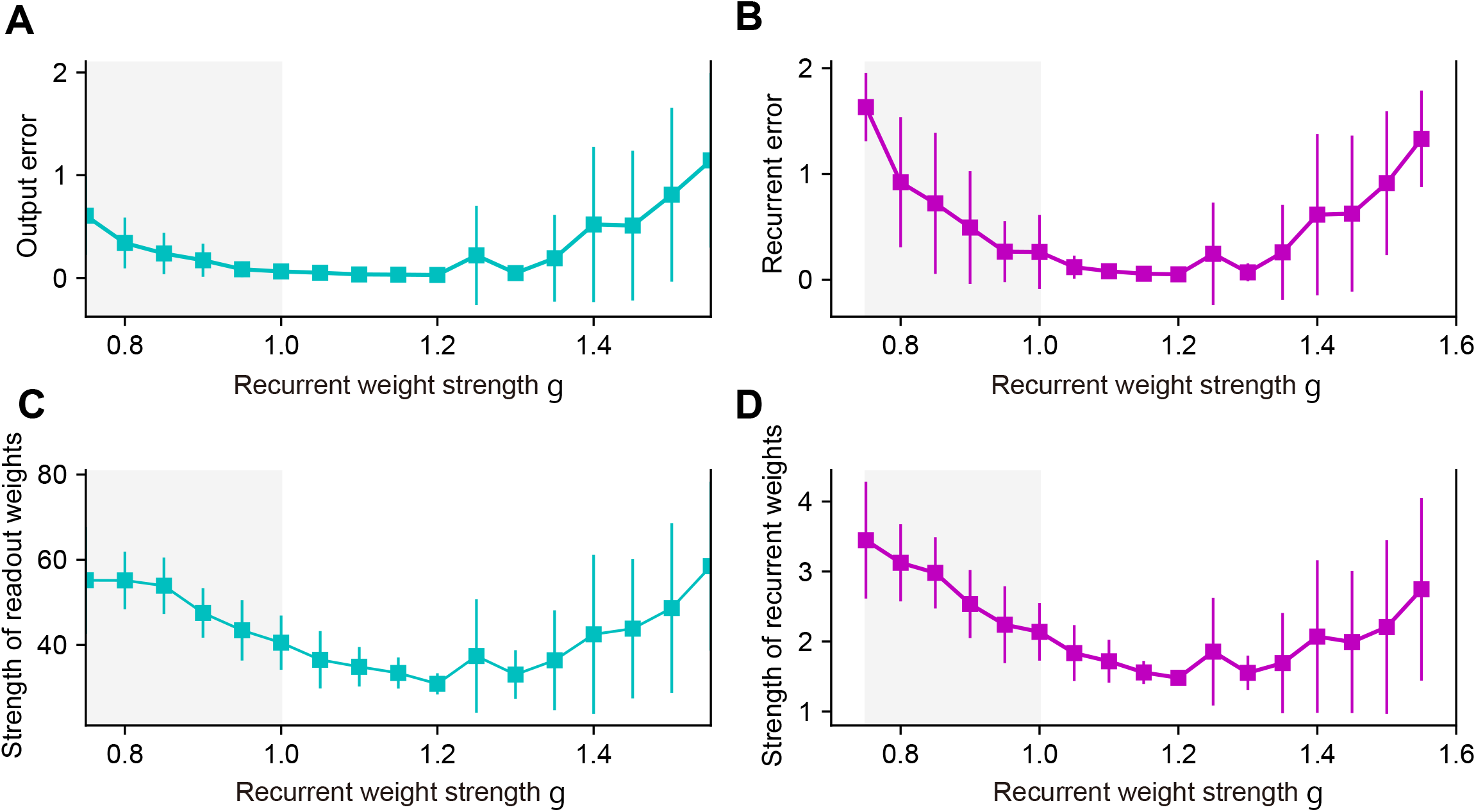
Crucial role of edge of chaos. (A) Output error of the trained network over various varue of recurrent strength. (B) Same as in A, but for the recurrent error. (C) Strength of trained readout weights are shown over various varue of recurrent strength. (D) Same as in C, but for the recurrent connection. Error bars stand for s.d.s over 20 independent simulations. In all figures, g=1 indicates that the network is on the edge of chaos.

**Supplementary Figure 4.**
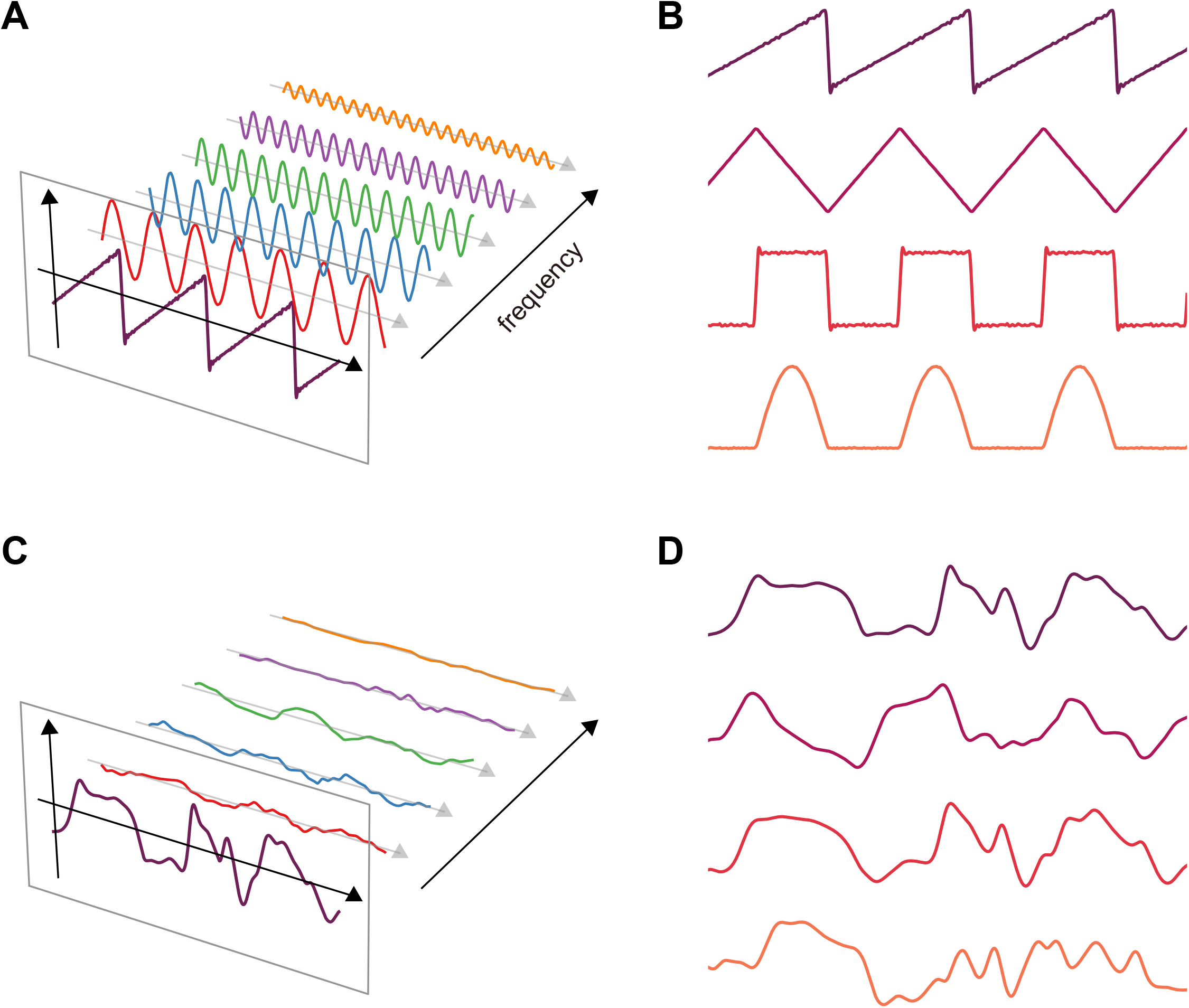
Generalization of learned representations. (A) The readout units were divided into two groups, and the training phase consisted of two stages. During the first stage of learning, readouts in the first group were trained to generate sinusoids with multiple frequencies (colored sinusoids). All readout weights projecting to the first group of readouts and the recurrent connections were trained in the first stage. In the second stage of learning, only the readout weights projecting to the second goup of readouts were trained to generate complex signals (purple discontinuous signal). (B)The model generalized multiple sinusoids to various complex signals simultaneously. (C) Same as in A, but without initial training. (D) Same as in B, but without initial training.

